# Synonymous mutations and the molecular evolution of SARS-Cov-2 origins

**DOI:** 10.1101/2020.04.20.052019

**Authors:** Hongru Wang, Lenore Pipes, Rasmus Nielsen

## Abstract

Human severe acute respiratory syndrome coronavirus 2 (SARS-CoV-2) is most closely related, by average genetic distance, to two coronaviruses isolated from bats, RaTG13 and RmYN02. However, there is a segment of high amino acid similarity between human SARS-CoV-2 and a pangolin isolated strain, GD410721, in the receptor binding domain (RBD) of the spike protein, a pattern that can be caused by either recombination or by convergent amino acid evolution driven by natural selection. We perform a detailed analysis of the synonymous divergence, which is less likely to be affected by selection than amino acid divergence, between human SARS-CoV-2 and related strains. We show that the synonymous divergence between the bat derived viruses and SARS-CoV-2 is larger than between GD410721 and SARS-CoV-2 in the RBD, providing strong additional support for the recombination hypothesis. However, the synonymous divergence between pangolin strain and SARS-CoV-2 is also relatively high, which is not consistent with a recent recombination between them, instead it suggests a recombination into RaTG13. We also find a 14-fold increase in the *d*_*N*_/*d*_*S*_ ratio from the lineage leading to SARS-CoV-2 to the strains of the current pandemic, suggesting that the vast majority of non-synonymous mutations currently segregating within the human strains have a negative impact on viral fitness. Finally, we estimate that the time to the most recent common ancestor of SARS-CoV-2 and RaTG13 or RmYN02 based on synonymous divergence, is 51.71 years (95% C.I., 28.11-75.31) and 37.02 years (95% C.I., 18.19-55.85), respectively.

## Introduction

The Covid19 pandemic is perhaps the biggest public health and economic threat that the world has faced for decades (Li, Guan, et al. 2020; Wu, et al. 2020; Zhou, Yang, et al. 2020). It is caused by a coronavirus (Lu, et al. 2020; Zhang and Holmes 2020), Severe acute respiratory syndrome coronavirus 2 (SARS-CoV-2), an RNA virus with a 29,903 bp genome consisting of four major structural genes (Wu, et al. 2020; Zhou, Yang, et al. 2020). Of particular relevance to this study is the *spike* protein which is responsible for binding to the primary receptor for the virus, angiotensin-converting enzyme 2 *(ACE2)* (Wan, et al. 2020; Wu, et al. 2020; Zhou, Yang, et al. 2020).

Human SARS-CoV-2 is related to a coronavirus (RaTG13) isolated from the bat *Rhinolophus affinis* from Yunnan province of China (Zhou, Yang, et al. 2020). RaTG13 and the human strain reference sequence (Genbank accession number MN996532) are 96.2% identical and it was first argued that, throughout the genome, RaTG13 is the closest relative to human SARS-CoV-2 (Zhou, Yang, et al. 2020). And RaTG13 and SARS-CoV-2 were 91.02% and 90.55% identical, respectively, to coronaviruses isolated from Malayan pangolins (Pangolin-CoV) seized at the Guangdong customs of China, which therefore form a close outgroup to the SARS-CoV-2+RaTG13 clade (Zhang, et al. 2020). Furthermore, five key amino acids in the receptor-binding domain (RBD) of *spike* were identical between SARS-CoV-2 and Pangolin-CoV, but differed between those two strains and RaTG13 (Zhang, et al. 2020). Xiao et al assembled and analysed a full-length Pangolin-CoV genome sequence, showing that the receptor-binding domain of its S protein differs from the SARS-CoV-2 by only one noncritical amino acid (Xiao, et al. 2020). Similar observations were made using Pangolin-CoV strains found in Malayan pangolin samples seized by the Guangxi customs of China (Lam, et al. 2020). Additionally, it is shown that when analyzing a window of length 582bp in the RBD, nonsynonymous mutations support a phylogenetic tree with SARS-CoV-2 and Pangolin-CoV as sister-groups, while synonymous mutations do not (Lam, et al. 2020). They discuss two possible explanations for their results, one which includes recombination and another which includes selection-driven convergent evolution. Independent analysis also support SARS-CoV-2 obtains the receptor binding motif through recombination from a donor related to this Pangolin-CoV strain (Li, Giorgi, et al. 2020). Detailed phylogenetic analysis on sub-regions across the S protein showed that it is the RaTG13 sequence that show exceptionally divergent pattern in the RBD region, they instead argued a recombination occurred into RaTG13 from an unknown divergent source (Boni, et al. 2020). This would explain the amino acid similarity between SARS-CoV-2 and Pangolin-CoV in the RBD as an ancestral trait that has been lost (by recombination) in RaTG13. Using a phylogenetic analysis they also dated the RaTG13 and SARS-CoV-2 divergence to be between 40 to 70 years. Recently, Zhou et al. discovered a viral strain, RmYN02 from the bat *Rhinolophus malayanus,* with a reported 97.2% identity in the ORF1ab gene but with only 61.3% sequence similarity to SARS-CoV-2 in the RBD (Zhou, Chen, et al. 2020). Moreover, the RmYN02 strain also harbors multiple amino acid insertions at the S1/S2 cleavage site in the spike protein (Zhou, Chen, et al. 2020).

To analyze the history of these sequences further, we here focus on patterns of synonymous divergence, which has received less focus, but also is less likely to be affected by selection than amino acid divergence. We develop a bias corrected estimator of synonymous divergence specific for SARS-CoV-2 and related strains, and analyze divergence using both sliding windows and a whole-genome approach between SARS-CoV-2 and related viral strains.

## Materials and methods

### BLAST searches

Sequences for blast databases were downloaded on March 26, 2020 from the following sources: EMBL nucleotide libraries for virus (ftp://ftp.ebi.ac.uk/pub/databases/embl/release/std), NCBI Virus Genomes (ftp://ftp.ncbi.nlm.nih.gov/genomes/Viruses), NCBI Virus Genbank Entries (ftp://ftp.ncbi.nlm.nih.gov/genomes/genbank/viral/), NCBI Influenza Genomes (ftp://ftp.ncbi.nlm.nih.gov/genomes/INFLUENZA/), all Whole Genome Shotgun (https://www.ncbi.nlm.nih.gov/genbank/wgs/) assemblies under taxonomy ID 10239, along with GISAID Epiflu and EpiCoV databases. Recently published sequences from the Myanmar bat samples (Valitutto, et al. 2020) were also added to the database. Blast databases were created using the default parameters for makeblastdb. Blast searches were performed using blastn (Altschul, et al. 1990) with parameters “-word_size 7 -reward 1 -penalty -3” and all other parameters as the default settings. All the blast hits to different Guangdong pangolin viral strain sequences were merged as one hit, and the blast hits to different Guangxi pangolin viral strain sequences were also merged.

### Alignment

To obtain an in-frame alignment of the genomes, we first identified the coding sequences of each viral strain using independent pairwise alignments with the coding sequences of the SARS-CoV-2 (Wuhan-Hu-1) genome. The genome alignments were performed using MAFFT (v7.450) (Katoh and Standley 2013) with parameters “--maxiterate 1000 --localpair”. The coding sequences of each gene were aligned using PRANK (Loytynoja 2014) (v.170427) with parameters “-codon -F”. Finally, the alignments for all genes were concatenated following their genomic order. ORF1a was excluded since its sequence is a subset of ORF1ab.

### Recombination detection

We detected possible recombination events across the genome using a combination of 7 alogorithms, RDP (Martin and Rybicki 2000), GENECONV (Padidam, et al. 1999), Bootscan (Salminen, et al. 1995), Maxchi (Smith 1992), Chimaera (Posada and Crandall 2001), SiSscan (Gibbs, et al. 2000), and 3Seq (Boni, et al. 2007) implemented in RDP5 program (Martin, et al. 2015) (version Beta 5.5) and then considered the recombination signals that were supported by at least two methods. We note that these 7 methods are all based on inferring recombination using the same type of evidence, and concordance between the methods cannot be interpreted as validation of the recombination signal. However, we will also use phylogenetic methods and methods based on relative sequence divergence to further investigate the putative recombination signals. The analysis was performed on the multiple sequence alignment consisting of the five viral strains. All regions showing recombination signals (Supplementary Table 5) were removed in subsequent analyses from all strains when stating that recombination regions were removed.

### Tree estimation

We estimated phylogenetic trees using two methods: Neighbor Joining (NJ) and Maximum Likelihood (ML). The NJ trees were estimated using *d*_*N*_ or *d*_*S*_ distance matrices which estimated using codeml (Yang 2007) with parameters “ runmode= -2, CodonFreq = 2, cleandata = 1“. To obtain bootstrap values, we bootstrapped the multiple sequence alignments 1,000 times, repeating the inference procedure for each bootstrap sample. The NJ tree was estimated using the ’neighbor’ software from the PHYLIP package (Felsenstein 2009). For ML trees, we used IQ-TREE (Nguyen, et al. 2015) (v1.5.2) with parameter “-TEST -alrt 1000” which did substitution model selection for the alignments and performed maximum-likelihood tree estimation with the selected substitution model for 1,000 bootstrap replicates. For this analysis, we masked all regions (Supplementary Table 5) that show recombination signals in any of the five studied viral genome. We masked regions from all sequences when at least one sequence showed evidence for recombination in that region. All masked regions are listed in Supplementary Table 5. The coordinates (based on the Wuhan-Hu-1 genome) of the three recombination regions (merged set of all the regions in Supplementary Table 5) were: 14611­15225, 21225-24252 and 25965-28297. We also estimate genome-wide divergence between RaTG13 and Wuhan-Hu-1 only excluding the region (position 22853-23092) where potential recombination was detected for the Wuhan-Hu-1 strain (Supplementary Table 5).

### Simulations

We simulated divergence with realistic parameters for SARS-CoV-2 using a continuous time Markov chain under the F3×4 codon-based model (Goldman and Yang 1994; Muse and Gaut 1994) (Yang, et al. 2000), which predicts codon frequencies from the empirical nucleotide frequencies in all 3 codon positions and using the global genomic maximum likelihood estimates of the transition/transversion bias *k(* =2.9024) and the *d_N_/d_S_* ratio ω (=0.0392) estimated from the human SARS-CoV-2 comparison to the nearest outgroup sequence, RaTG13 (see Results). For the simulations of short 300 bp sequences we kept *ω* constant but varied time such that the number of synynoymous substitutions per synonymous sites, *d*_*S*_, varied between 0.25 and 3.00. Estimates of *d*_*S*_ > 3 are truncated to 3. For simulations of genome-wide divergence between RaTG13 and human strains, we fix *d*_*S*_ at 0.1609 (the maximum likelihood estimate outside the RBD region reported in the Results section). In all cases, we use 10,000 independent replicate simulations for each parameter setting.

### Estimation of sequence divergence in 300-bp windows

*d*_*N*_ and *d*_*S*_ were estimated using two different methods implemented in the PAML package (Yang 2007) (version 4.9d): a count-based method, YN00 (Yang and Nielsen 2000) as implemented in the program ‘yn00’ with parameters “icode = 0, weighting = 0, commonf3×4 = 0”, and a maximum-likelihood method (Goldman and Yang 1994; Muse and Gaut 1994) implemented in codeml applied with arguments “runmode= -2, CodonFreq = 2”. The estimates in 300-bp windows were further bias-corrected as described below.

### Bias correction for d_S_ estimates in 300-bp window

To correct for the biases observed in the estimation of *d*_*S*_ (see results section) we identifed a quartic function which maps from 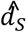, the estimates of *d*_*S*_, into 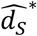, the bias corrected estimate such that to a close approximation, 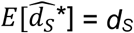. To identify the coefficients of this function we used 10,000 simulations as previously described, on a grid of *d*_*S*_ values (0.25, 0.5, 0.75, …, 3.0). We then identified coefficients such that sum of 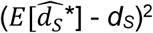 is minimized over all simulation values.

## Results

### Database searches

The genome of human coronavirus can effectively recombine with other viruses to form a chimeric new strain when they co-infect the same host (Forni, et al. 2017; Boni, et al. 2020). Complicated recombination histories have been observed in the receptor binding motif region of the spike protein (Lam, et al. 2020; Xiao, et al. 2020; Zhang, et al. 2020) and several other regions (Boni, et al. 2020) of the SARS-CoV-2, it is thus important to exhaustively search along the viral genome for other regions potentially of recombination origin and identify possible donors associated with them. To identify possible viral strains that may have contributed, by recombination, to the formation of human SARS-CoV-2, we searched NCBI and EMBL virus entries along with GISAID Epiflu and EpiCov databases for similar sequences using BLAST in 100bp windows stepping every 10bp (Fig. 1b). The majority of the genome (78.1%, 2330/2982 of the windows) has one unique best hit, likely reflecting the high genetic diversity of the coronavirus. 21.9% of the genomic regions has multiple best hits, which suggests that these regions might be more conserved. Among the windows with unique best hits, 97.0% (2260/2330) of them were the RaTG13 or RmYN02 bat strains and 1.9% of them, including the *ACE2* contact residues region of the S protein, were the pangolin SARS-CoV-2 virus. These observations are consistent with previous results that RaTG13 and RmYN02 are the most closely related viral strains, while the region containing the *ACE2* contact residues is more closely related to the pangolin virus strain (Lam, et al. 2020; Li, Giorgi, et al. 2020; Xiao, et al. 2020; Zhang, et al. 2020). A considerable amount of genomic regions (20 windows with unique hits) show highest sequence identity with other coronaviruses of the SARS-CoV-2 related lineage (Lam, et al. 2020) (bat-SL-CoVZC45 and bat-SL-CoVZXC21 (Hu, et al. 2018)). In addition, there were 6 windows whose unique top hits are coronavirus of a SARS-CoV related lineage (Lam, et al. 2020) (Supplementary Table 4). The mosaic pattern that different regions of the genome show highest identity to different virus strains is likely to have been caused by the rich recombination history of the SARS-CoV-2 lineage (Boni, et al. 2020; Li, Giorgi, et al. 2020; Patino-Galindo, et al. 2020). Moreover, its unique connection with SARS-CoV related lineages in some genomic regions may suggest recombination between the ancestral lineage of SARS-CoV-2 and distantly related virus lineages, although more formal analyses are needed to determine the recombination history (see also Boni, et al. 2020 for further discussion).

**Figure 1.**
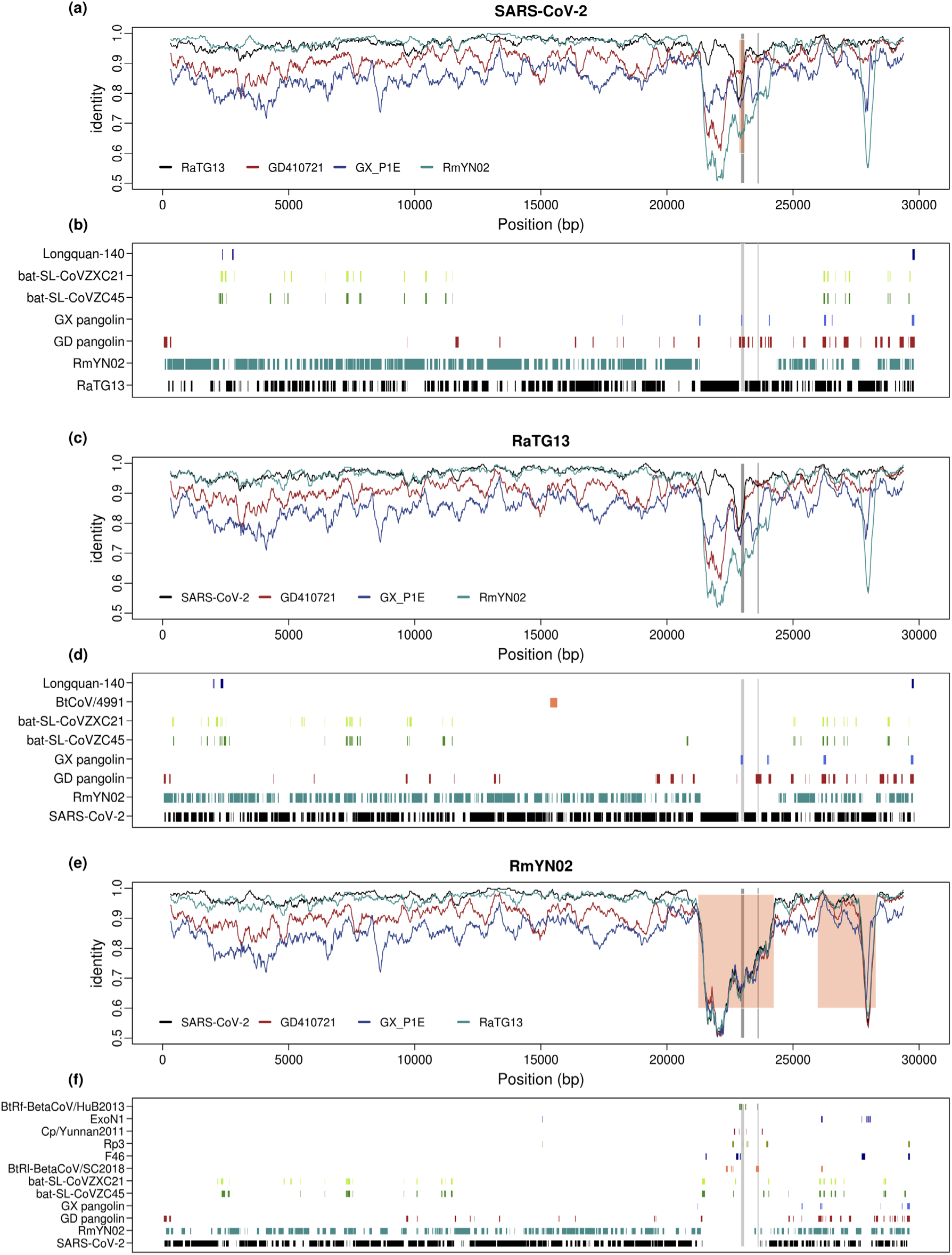
Genome-wide identity plot and top blast hits for SARS-CoV-2, RaTG13 and RmYN02. (a) 300 bp sliding-windows of nucleotide identity between SARS-CoV-2 and the four most closely related viral strains, RmYN02, RaTG13, GD410721 and GX_P1E. Orange shading marks the recombinant region in SARS-CoV-2 inferred by 3SEQ (details in Supplementary Table 5). (b) the plot lists all the viral strains that are the unique best BLAST hit in at least three 100-bp windows, when blasting with SARS-CoV-2, with the regions where each strain is the top blast hit marked. (b) and (c). Figures for RaTG13 (c, d) and RmYN02 (e, f) generated in the same way as for SARS-CoV-2 in (a) and (b). The *ACE2* contact residues of RBD region (left) and the furin sites (right) of the S protein are marked in both plots with grey lines.

Searching databases with BLAST using the most closely related viral strains, RaTG13 and RmYN02, we observe a very similar pattern, as that observed for SARS-CoV-2, in terms of top hits across the genome (Fig. 1b), suggesting that these possible recombination events with distantly related lineages are not unique to the SARS-CoV-2 lineage, but happened on the ancestral lineage of SARS-CoV-2, RaTG13, and RmYN02. A notable exception is a large region around the S gene, where RmYN02 show little similarity to both SARS-CoV-2 and RaTG13.

### Sequence similarity and recombination

We focus further on studying the synonymous evolution of SARS-CoV-2, and analyzing Wuhan-Hu-1 as the human nCoV19 reference strain (Wu, et al. 2020) along with the four viral strains with highest overall identity: the bat strains RmYN02 and RaTG13 (Zhou, Chen, et al. 2020; Zhou, Yang, et al. 2020), and the Malayan pangolin strains, GD410721 and GX_P1E, which were isolated from Malayan pangolin samples seized by Guangdong and Guangxi Customs of China, respectively. These four strains have previously been identified as the strains most closely related to SARS-CoV-2 (Lam, et al. 2020; Xiao, et al. 2020). Other available phylogenetically related, but less similar viral strains, such as bat-SL-CoVZXC21 and bat-SL-CoVZC45 (Hu, et al. 2018), are not included due to nearly saturated synonymous mutations when compared with SARS-CoV-2 (maximum likelihood estimates of *d*_*S*_ = 3.2067 and 2.8445, respectively).

We performed recombination analyses across the five viral genomes based on the concensus of the seven recombination-detection methods implemented in RDP5 (see Methods). We identified nine recombination regions affecting at least one of the sequences (Supplementary Table 5). Phylogenetic analyses of these regions confirm phylogenetic incongruence when compared with genome-wide trees (Fig. 2 and Supplementary Figure 1–3). Particularly, a recombination signal is found in a region encompassing the RBD of the S protein, suggesting that the human SARS-CoV-2 (Wuhan-Hu-1) sequence is a recombinant with the Pangolin-CoV (GD410721) as the donor (Supplementary Table 5). Phylogenetic analyses also support that Wuhan-Hu-1 and GD410721 form a clade relative to RaTG13 (Supplementary Figure 1c, 1d). Phylogenetic analyses (Fig. 2) in genomic regions with all recombination tracts (Supplementary Table 5) masked using Maximum-likelihood (Fig. 2a) and Neighbor-joining based on synonymous (Fig. 2b) or non-synoymous (Fig. 2c) mutation distance metrics, consistently support RmYN02 as the nearest outgroup to human SARS-CoV-2, in contrast to previous analyses before the discovery of RmYN02, which instead found RaTG13 to be the nearest outgroup (Lam, et al. 2020; Wu, et al. 2020). This observation is also consistent with the genome-wide phylogeny constructed in previous study (Zhou, Chen, et al. 2020).

**Figure 2.**
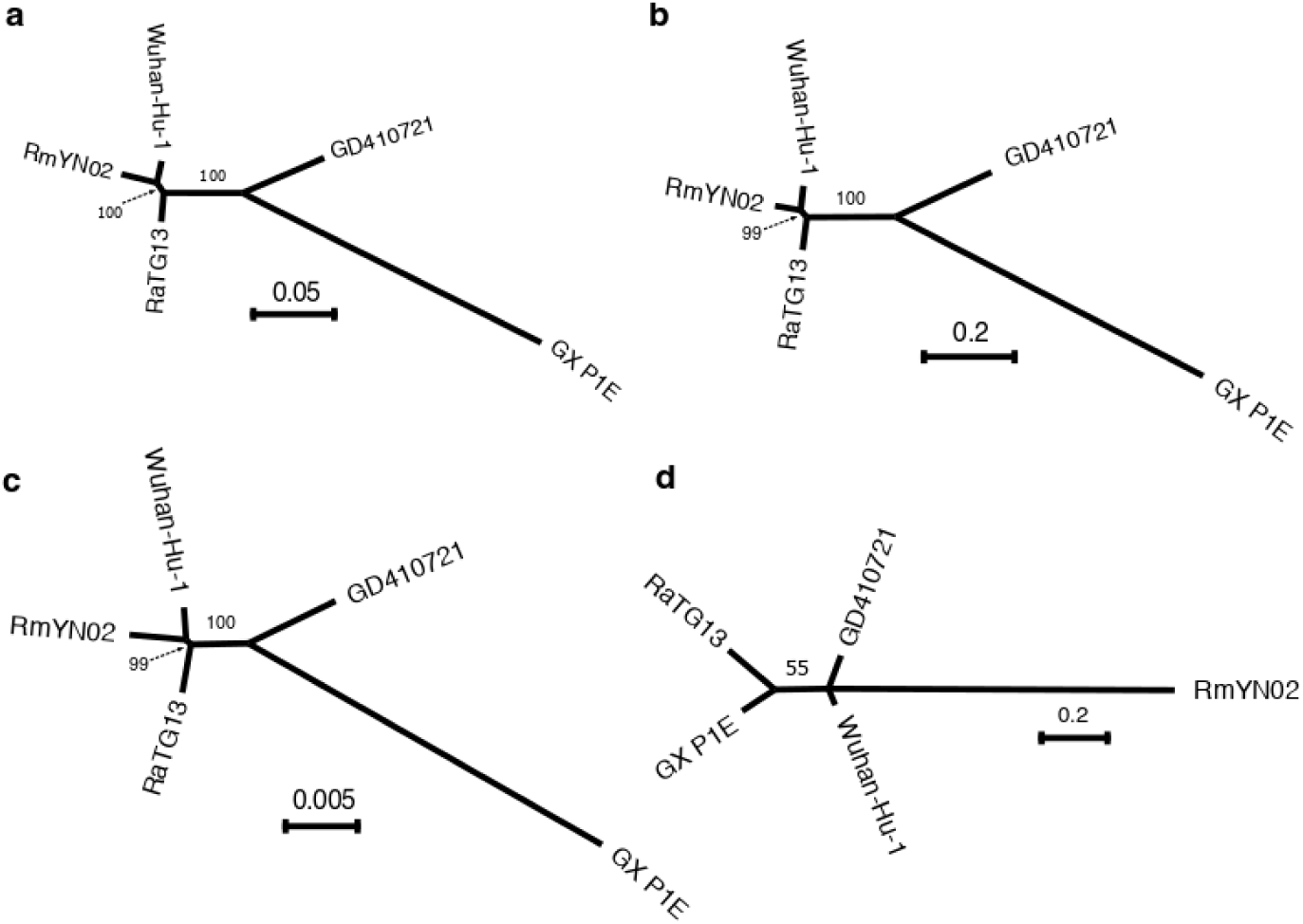
Unrooted phylogenies of the virus strains. (a) Maximum-likelihood tree in genomic regions with recombination tracts removed; (b) Neighbor-joining tree using synonymous mutation (*d*_*S*_) distance in genomic regions with recombination tracts removed; (c) Neighbor-joining tree using non-synonymous mutation (*d*_*N*_) distances in genomic regions with recombination tracts removed; (d) The maximum-likelihoods tree at the receptor-binding domain *ACE2* contact residues (51 amino acids) region. The bootstrap values are based on 1,000 replicates. The associated distance matrix for (b) and (c) can be found in Table 3.

We plot the overall sequence similarity (% nucleotides identical) between SARS-CoV-2 and the four other strains analyzed in windows of 300 bp (Fig. 1). Notice that the divergences between human SARS-CoV-2 and the bat viral sequences, RaTG13 and RmYN02, in most regions of the genome, are quite low compared to the other comparisons. A notable exception is the suspected recombination region in RmYN02 that has an unusual high level of divergence with all other viruses (Fig. 1e). However, there is also another exception: a narrow window in the RBD of the S gene where the divergence between SARS-CoV-2 and GD410721 is moderate and the divergences between GD410721 and both SARS-CoV-2 and RaTG13 are quite high and show very similar pattern. This, as also found in the recombination analyses based on methdos implemented in RDP5, would suggest a recombination event from a strain related to GD410721 into an ancestor of the human strain (Lam, et al. 2020; Xiao, et al. 2020; Zhang, et al. 2020), or alternatively, from some other species into RaTG13, as previously hypothesized (Boni, et al. 2020). We note that RmYN02 is not informative about the nature of this event as it harbors a long and divergent haplotype in this region, possibly associated with another independent recombination event with more distantly related viral strains (Fig. 1e). The other four sequences are all highly, and approximately equally, divergent from RmYN02 in this large region (Fig. 1e), suggesting that the RmYN02 strain obtained a divergent haplotype from the recombination event. When BLAST searching using 100-bp windows along the RmYN02 genome, we find no single viral genome as the top hit, instead the top hits are found sporadically in different viral strains of the SARS-CoV lineage (Fig. 1f), suggesting that the sequence of the most proximal donor is not represented in the database.

### Estimating synonymous divergence and bias correction

While the overall divergence in the S gene encoding the *spike* protein could suggest the presence of recombination in the region, previous study (Lam, et al. 2020) reported that the tree based on synonymous substitutions supported RaTG13 as the sister taxon to the human SARS-CoV-2 also in this region. That would suggest the similarity between GD410721 and human SARS-CoV-2 might be a consequence of convergent evolution, possibly because both strains adapted to the use of the same receptor. An objective of the current study is to examine if there are more narrow regions of the spike protein that might show evidence of recombination. We investigate this issue using estimates of synonymous divergence per synonymous site (*d*_*S*_) in sliding windows of 300 bp. However, estimation of *d*_*S*_ is complicated by the high levels of divergence and extremely skewed nucleotide content in the 3rd position of the sequences (Table 1) which will cause a high degree of homoplasy. We, therefore, entertain methods for estimation that explicitly account for unequal nucleotide content and multiple hits in the same site such as maximum likelihood methods and the YN00 method (Yang and Nielsen 2000). It is shown that for short sequences, some counting methods, such as the YN00 method, can perform better in terms of Mean Squared Error (MSE) for estimating *d*_*N*_ and *d*_*S*_ (Yang and Nielsen 2000). However, it is unclear in the current case how best to estimate *d*_*S*_. For this reason, we performed a small simulations study (see Methods) for evaluating the performance of the maximum likelihood (ML) estimator of *d*_*N*_ and *d*_*S*_ (as implemented in codeml (Yang 2007)) under the F3×4 model and the YN00 method implemented in PAML. In general, we find that estimates under the YN00 are more biased with slightly higher MSE than the ML estimate for values in the most relevant regime of *d*_*S*_ < 1.5 (Fig. 3). However, we also notice that both estimators are biased under these conditions. For this reason, we perform a bias correction calibrated using simulations specific to the nucleotide frequencies and *d_N_/*d_*S*_ ratio observed for SARS-CoV-2 (see Methods). The bias corrections we obtain are 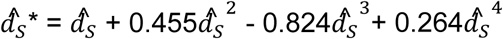, for the ML estimator and 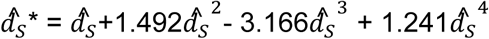 for yn00. Notice that there is a trade-off between mean and variance (Fig. 3) so that the MSE becomes very large, particularly for the for yn00 method, after bias correction. For *d*_*S*_ >2 the estimates are generally not reliable, however, we note that for *d*_*S*_ <1.5 the bias-corrected ML estimator tends overall to have slightly lower MSE, and we, therefore, use this estimator for analyses of 300 bp regions.

**Figure 3.**
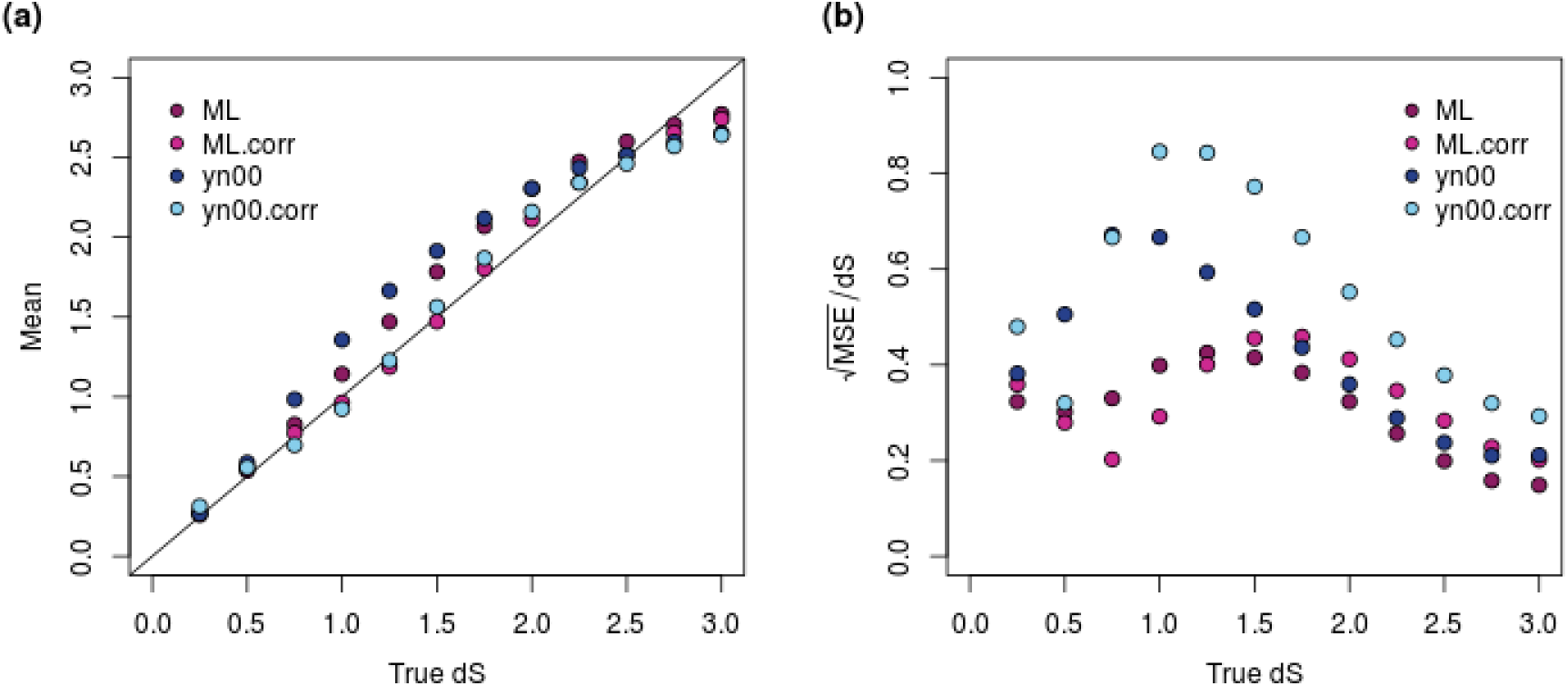
Bias correction for *d*_*S*_ estimate in 300-bp windows. (a) The mean of *d*_*S*_ estimates using different methods; ML.corr and yn00.corr are the bias corrected versions of the ML and yn00 methods, respectively. (b) Errors in *d*_*S*_ estimates as measured using the ratio of square root of mean squared error (MSE) to true *d*_*S*_. All the estimates are based on 10,000 simulations. ML: maximum-likelihood estimates using the f3×4 model in codeml; ML.corr, maximum-likelihood estimates with bias correction; yn00, count-based estimates in (Yang and Nielsen 1999); yn00.corr, yn00 estimates with bias correction. All *d*_*S*_ estimates are truncated at 3, explaining the reduction in MSE with increasing values of *d*_*S*_ as *d*_*S*_ approaches 3.

**Table 1.**
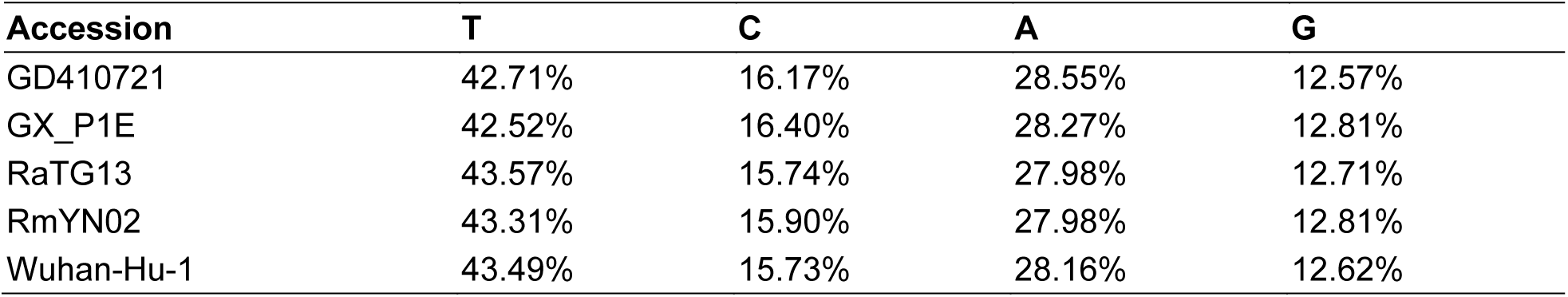
Genome-wide nucleotide composition at the third position of the codons in the viral strains. The nucletodie compositions at the first and second positions can be found in Supplementary table 1.

| Accession | T | C | A | G |
| --- | --- | --- | --- | --- |
| GD410721 | 42.71% | 16.17% | 28.55% | 12.57% |
| GX_P1E | 42.52% | 16.40% | 28.27% | 12.81% |
| RaTG13 | 43.57% | 15.74% | 27.98% | 12.71% |
| RmYN02 | 43.31% | 15.90% | 27.98% | 12.81% |
| Wuhan-Hu-1 | 43.49% | 15.73% | 28.16% | 12.62% |

### Synonymous divergence

We estimate *d*_*N*_ and *d*_*S*_ under the F3×4 model in codeml (Goldman and Yang 1994; Muse and Gaut 1994) and find genome-wide estimates of *d*_*S*_ = 0.1604, *d*_*N*_ = 0.0065 *(d*_*N*_*/ d_S_* = 0.0405) between SARS-CoV-2 and RaTG13 and *d*_*S*_ = 0.2043, *d*_*N*_ = 0.0220 *(d_N_/d_S_* = 0.1077) between SARS-CoV-2 and RmYN02. However, a substantial amount of this divergence might be caused by recombination with more divergent strains. We, therefore, also estimate *d*_*N*_ and *d*_*S*_ for the regions with inferred recombination tracts (Supplementary Table 5) removed from all sequences (Table 3). We then find values of *d_S_ =* 0.1462 (95% C.I., 0.1340-0.1584) and *d*_*S*_ = 0.1117 (95% C.I., 0.1019-0.1215) between SARS-CoV-2 and RaTG13 and RmYN02, respectively. This confirms that RmYN02 is the virus most closely related to SARS-CoV-2. The relative high synonymous divergence also shows that the apparent high nucleotide similarity between SARS-CoV-2 and the bat strains (96.2% (Zhou, Yang, et al. 2020) and 97.2%(Zhou, Chen, et al. 2020)) is caused by conservation at the amino acid level *(d*_*N*_*/ d*_*S*_ = 0.0410 and 0.0555) exacerbated by a high degree of synonymous homoplasy facilitated by a highly skewed nucleotide composition at the third position of codons (with an AT content >72%, Table 1).

The synonymous divergence to the pangolin sequences GD410721 and GX_P1E in genomic regions with inferred recombination tracts removed is 0.5095 (95% C.I., 0.4794-0.5396) and 1.0304 (95% C.I., 0.9669-1.0939), respectively. Values for other comparisons are shown in Tables 2 and 3. In comparisons between SARS-CoV-2 and more distantly related strains, *d*_*S*_ will be larger than 1, and with this level of saturation, estimation of divergence is associated with high variance and may be highly dependent on the accuracy of the model assumptions. This makes phylogenetic analyses based on synonymous mutations unreliable when applied to these more divergent sequences. Nonetheless, the synonymous divergence levels seem generally quite compatible with a molecular clock with a *d*_*S*_ of 0.9974 (95% C.I., 0.9381-1.0567, GD410721), 1.0366 (95% C.I., 0.9737-1.0995, RaTG13), 1.0333 (95% C.I., 0.9699-1.0967, RmYN02) and 1.0304 (95% C.I., 0.9669-1.0939, Wuhan-Hu-1) between the outgroup, GX_P1E, and the three ingroup strains. The largest value is observed for RaTG13 *(d*_*S*_ = 1.0366), despite this sequence being the most early sampled sequence, perhaps caused by additional undetected recombination into RaTG13.

**Table 2.** Whole genome *d*_*N*_ and *d*_*S*_ estimates among the viral strains. The *d*_*S*_ estimates are shaded in green, and the *d*_*N*_ estimates are in orange shade. The 95% confidence intervals, calculated based on 1,000 bootstrap replicates, are included in the brackets for each estimates.

|  | GD410721 | GX_P1E | RaTG13 | RmYN02 | Wuhan-Hu-1 |
| --- | --- | --- | --- | --- | --- |
| GD410721 |  | 0.0372<br>(0.0341-0.0403) | 0.0171<br>(0.0152-0.0190) | 0.0293<br>(0.0266-0.0320) | 0.0160<br>(0.0142-0.0178) |
| GX_P1E | 0.9883<br>(0.9338-1.0428) |  | 0.0347<br>(0.0318-0.0376) | 0.0485<br>(0.0450-0.0520) | 0.0342<br>(0.0314-0.0370) |
| RaTG13 | 0.5392<br>(0.5105-0.5679) | 1.0156<br>(0.9608-1.0704) |  | 0.0235<br>(0.0210-0.0260) | 0.0065<br>(0.0053-0.0077) |
| RmYN02 | 0.6001<br>(0.5681-0.6321) | 1.0757<br>(1.0166-1.1348) | 0.2438<br>(0.2285-0.2591) |  | 0.0220<br>(0.0195-0.0245) |
| Wuhan-Hu-1 | 0.5425<br>(0.5131-0.5719) | 0.9973<br>(0.9434-1.0512) | 0.1604<br>(0.1491-0.1717) | 0.2043<br>(0.1901-0.2185) |  |

**Table 3.** Genome-wide *d*_*N*_ and *d*_*S*_ estimates after removing recombination regions inferred by 3SEQ. The *d*_*S*_ estimates are shaded in green, and the *d*_*N*_ estimates are in orange shade. The coordinates relative to the Wuhan-Hu-1 genome of the masked region can be found in the method section. The 95% confidence intervals, calculated based on 1,000 bootstrap replicates, are included in the brackets for each estimates.

|  | GD410721 | GX_P1E | RaTG13 | RmYN02 | Wuhan-Hu-1 |
| --- | --- | --- | --- | --- | --- |
| GD410721 |  | 0.0348<br>(0.0317-0.0379) | 0.0138<br>(0.0120-0.0156) | 0.0152<br>(0.0133-0.0171) | 0.0135<br>(0.0117-0.0153) |
| GX_P1E | 0.9974<br>(0.9381-1.0567) |  | 0.0357<br>(0.0325-0.0389) | 0.0361<br>(0.0329-0.0393) | 0.0349<br>(0.0318-0.0380) |
| RaTG13 | 0.4962<br>(0.4669-0.5255) | 1.0366<br>(0.9737-1.0995) |  | 0.0079<br>(0.0066-0.0092) | 0.0060<br>(0.0048-0.0071) |
| RmYN02 | 0.5070<br>(0.4773-0.5366) | 1.0333<br>(0.9699-1.0967) | 0.1522<br>(0.1395-0.1649) |  | 0.0062<br>(0.0050-0.0074) |
| Wuhan-Hu-1 | 0.5095<br>(0.4794-0.5396) | 1.0304<br>(0.9669-1.0939) | 0.1462<br>(0.1340-0.1584) | 0.1117<br>(0.1019-0.1215) |  |

### Sliding windows of synonymous divergence

To address the issue of possible recombination we plot *d*_*S*_ between SARS-CoV-2, GD410721, and RaTG13 and the ratio of *d*_*S*_(SARS-CoV-2, GD410721) to *d*_*S*_(SARS-CoV-2, RaTG13) in 300 bp sliding windows along the genome. Notice that we truncate the estimate of *d*_*S*_ at 3.0. Differences between estimates larger than 2.0 should not be interpreted strongly, as these estimates have high variance and likely will be quite sensitive to the specifics of the model assumptions.

We find that *d*_*S*_(SARS-CoV-2, GD410721) approximately equals *d*_*S*_(GD410721, RaTG13) and is larger than *d*_*S*_(SARS-CoV-2, RaTG13) in almost the entire genome showing than in these parts of the genome GD410721 is a proper outgroup to (SARS-CoV-2, RaTG13) assuming a constant molecular clock. One noticeable exception from this is the RBD region of the S gene. In this region the divergence between SARS-CoV-2 and GD410721 is substantially lower than between GD410721 and RaTG13 (Fig. 4a,4c). The same region also has much smaller divergence between SARS-CoV-2 and GD410721 than between SARS-CoV-2 and RaTG13 (Fig. 4a,4c). The pattern is quite different than that observed in the rest of the genome, most easily seen by considering the ratio of *d*_*S*_(SARS-CoV-2, GD410721) to *d*_*S*_(SARS-CoV-2, RaTG13) (Fig. 2b, 2d). In fact, the estimates of *d*_*S*_(SARS-CoV-2, RaTG13) are saturated in this region, even though they are substantially lower than 1 in the rest of the genome. This strongly suggests a recombination event in the region and provides independent evidence of that previously reported based on amino acid divergence (e.g.,(Zhang, et al. 2020)).

**Figure 4.**
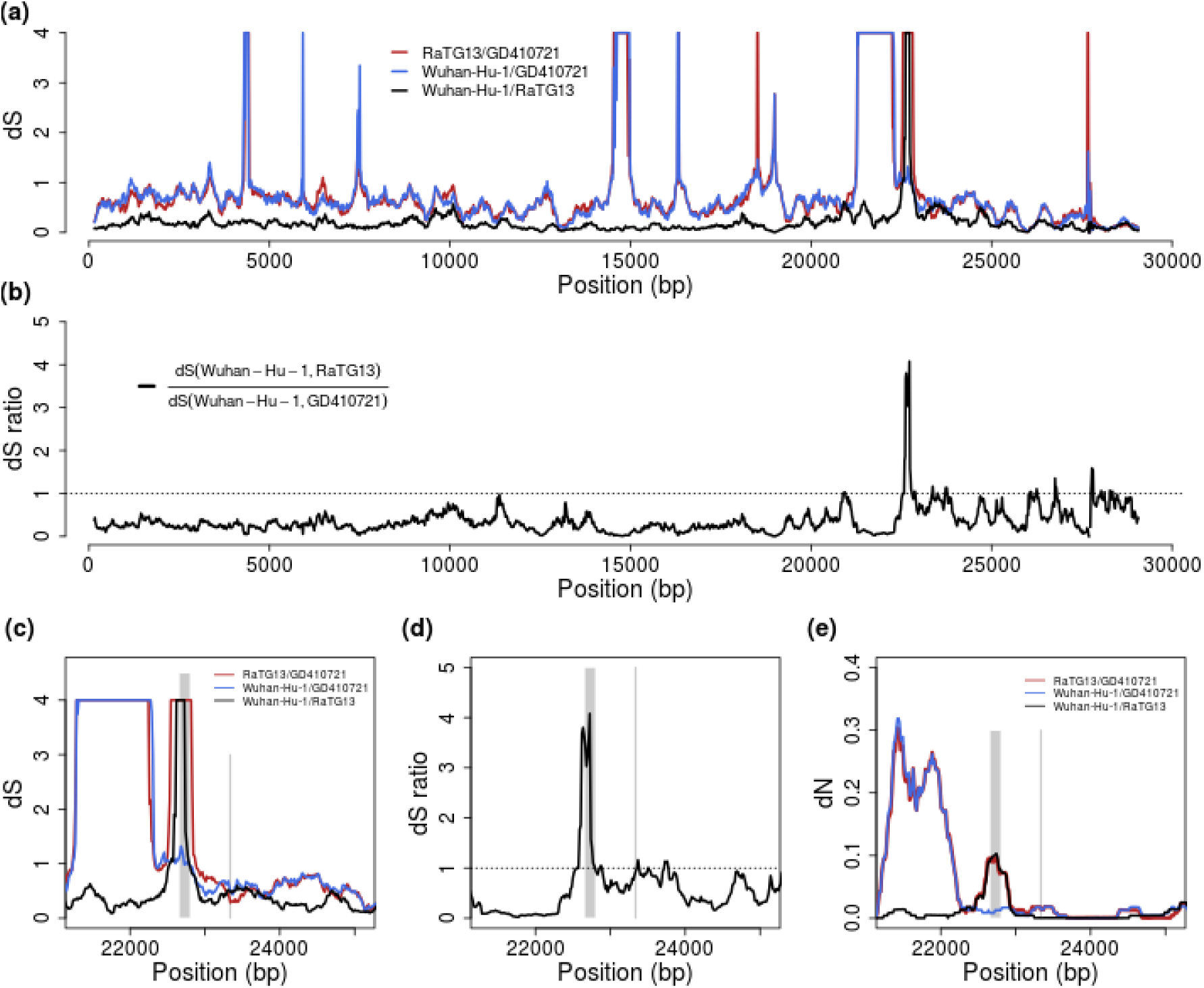
*d*_*S*_ and *d*_*N*_ estimates across the virus genome. (a) Pairwise *d*_*S*_ estimates in 300-bp sliding windows for RaTG13, GD410721 and Wuhan-Hu-1, the estimates are truncated at 4. (b) *d*_*S*_ ratio of *d*_*S*_ (Wuhan-Hu-1,RaTG13) to *d*_*S*_ (Wuhan-Hu-1,GD410721). (c) and (d) are the zoom-in plot for *d*_*S*_ and *d*_*S*_-ratio at the *spike* (S) protein region. The receptor-binding domain contact residues (left) and furin site regions (right) are marked with grey lines. (e) the pairwise *d*_*N*_ estimates in 300-bp sliding windows in the S protein for these strains. The *d*_*S*_ values are truncated at 4 in the plots.

The combined evidences from synonymous divergence and the topological recombination inference, provide strong support for the recombination hypothesis. However, these analyses alone do not distinguish between recombination into RaTG13 from an unknown source as previously hypothesized (Boni, et al. 2020) and recombination between SARS-CoV-2 and GD410721 as proposed as one possible explanation by Lam et al. (Lam, et al. 2020). To distinguish between these hypotheses we searched for sequences that might be more closely related, in the RBD region, to RaTG13 than SARS-CoV-2 and we plotted sliding window similarities across the genome for RaTG13 (Fig. 1c). We observe relatively low sequence identity between RaTG13 and all three other strains in the *ACE2* contact residue region of the *spike* protein, which is more consistent with the hypothesis of recombination into RaTG13, as proposed in (Boni, et al. 2020). Moreover, our BLAST search analyses of RaTG13 in this region show highest local sequence similarity with GX pangolin virus strains which is the genome-wide outgroup for the three other sequences (Lam, et al. 2020). This observation is more compatible with the hypothesis of recombination from a virus related to GX pangolin strains, than with recombination between SARS-CoV-2 and GD410721.

Unfortunately, because of the high level of synonymous divergence to the nearest outgroup, tree estimation in small windows is extremely labile in this region. In fact, synonymous divergence appears fully saturated in the comparison with GX_P1E, eliminating the possibility to infer meaningful trees based on synonymous divergence. However, we can use the overall maximum likelihood tree using both synonymous and nonsynonymous mutations (Fig. 2d). The ML tree using sequence from the *ACE2* contact residue region supports the clustering of SARS-CoV-2 and GD410721, but with unusual long external branches for all strains except SARS-CoV-2, possibly reflecting smaller recombination regions within the *ACE2* contact residue region.

### Weakly deleterious mutations and clock calibrations

The use of synonymous mutations provides an opportunity to calibrate the molecular clock without relying on amino acid changing mutations that are more likely to be affected by selection. The rate of substitution of weakly and slightly deleterious mutations is highly dependent on ecological factors and the effective population size. Weakly deleterious mutations are more likely to be observed over small time scales than over long time scales, as they are unlikely to persist in the population for a long time and go to fixation. This will lead to a decreasing *d*_*N*_*/d*_*S*_ ratio for longer evolutionary lineages. Furthermore, changes in effective population size will translate into changes in the rate of substitution of slightly deleterious mutations. Finally, changes in ecology (such as host shifts, host immune changes, changes in cell surface receptor, etc.) can lead to changes in the rate of amino acid substitution. For all of these reasons, the use of synonymous mutations, which are less likely to be the subject of selection than nonsynonymous mutations, are preferred in molecular clock calculations. For many viruses, the use of synonymous mutations to calibrate divergence times is not possible, as synonymous sites are fully saturated even at short divergence times. However, for the comparisons between SARS-CoV-2 and RaTG13, and SARS-CoV-2 and RmYN02, synonymous sites are not saturated and can be used for calibration. We find an estimate of ω = 0.0391 between SARS-CoV-2 and RaTG13, excluding just the small RDB region showing a recombination signal in SARS-CoV-2 (Supplementary Table 5, coordinates: 22851-23094). Using 1000 parametric simulations under the estimated values and the F3×4 codon model, we find that the estimate is approximately unbiased (ώ = 0.0398, S.E.M.= 0.0001) and with standard deviation 0.0033, providing an approximate 95% confidence interval of (0.0332, 0.0464). Also, using 59 human strains of SARS-CoV-2 from Genbank and National Microbiology Data Center (See Methods) we obtain an estimate of ω = 0.5604 using the F3×4 model in codeml. Notice that there is a 14­fold difference in *dN/d*_*S*_ ratio between these estimates. Assuming very little of this difference is caused by positive selection, this suggests that the vast majority of mutations currently segregating in the SARS-CoV-2 are slightly or weakly deleterious for the virus.

### Dating of divergence between Bat viruses and SARS-CoV-2

To calibrate the clock we use the estimate provided by (http://virological.org/t/phylodynamic-analysis-of-sars-cov-2-update-2020-03-06/420) of μ =1.04×10^−3^ substitutions/site/year (95% CI: 0.71×10-3, 1.40×10-3). The synonymous specific mutation rate can be found from this as *d*_*S*_/year = *μ*_*S*_ = *μ*_*l*_(*PS* + ω*pN*, where ω is the *d_N_/d*_*S*_ ratio, and *pN* and *pS* are the proportions of nonsynonymous and synonymous sites, respectively. The estimate of the total divergence on the two lineages is then 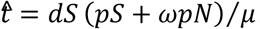 Inserting the numbers from Table 3 for the divergence between SARS-CoV-2 and RaTG13 and RmYN02 respectively, we find a total divergence of 96.92 years and 74.05 years respectively. Taking into account that RaTG13 was isolated July 2013, we find an estimated tMRCA between that strain and SARS-CoV-2 of 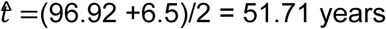. Similarly, we find an estimate of divergence between SARS-CoV-2 and RmYN02 of 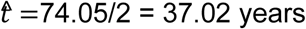, assuming approximately equal sampling times. The estimate for SARS-CoV-2 and RaTG13 is compatible with the values obtained using different methods for dating (Boni, et al. 2020). The variance in the estimate in *d*_*S*_ is small and the uncertainty is mostly dominated by the uncertainty in the estimate of the mutation rate. We estimate the S.D. in t using 1000 parametric simulations, using the ML estimates of all parameters, for both RaTG13 vs. SARS-CoV-2 and for RmYN02 vs. SARS-CoV-2, and for each simulated data also simulating values of *μ* and ω from normal distributions with mean 1.04×10^−3^ and S.D. 0.18×10^−3^, and mean 0.5604 and S.D. 0.1122, respectively. We subject each simulated data set to the same inference procedure as done on the real data. Our estimate of the S.D. in the estimate is 11.8 for RaTG13 vs. SARS-CoV-2 and 9.41 for RmYN02 vs. SARS-CoV-2, providing an approximate 95% confidence interval of (28.11, 75.31) and (18.19, 55.85), respectively. For RaTG13, if including all sites, except the 244-bp in the RBD of the S gene (Supplementary Table 5), the estimate is 55.02 years with an approx. 95% C.I. of (29.4, 80.7). As more SARS-CoV-2 sequences are being obtained, providing more precise estimates of the mutation rate, this confidence interval will become narrower. However, we warn that the estimate is based on a molecular clock assumption and that violations of this assumption eventually will become a more likely source of error than the statistical uncertainty quantified in the calculation of the confidence intervals. We also note that, so far, we have assumed no variation in the mutation rate among synonymous sites. However, just from the analysis of the 300 bp windows, it is clear that is not true. The variance in the estimate of *d*_*S*_ among 300 bp windows from the RaTG13-SARS-CoV-2 comparison is approximately 0.0113. In contrast, in the simulated data assuming constant mutation rate, the variance is approximately 0.0034, suggesting substantial variation in the synonymous mutation rate along the length of the genome. Alternatively, this might be explained by undetected recombination in the evolutionary history since the divergence of the strains.

## Discussion

The highly skewed distribution of nucleotide frequencies in synonymous sites in SARS-CoV-2 (Kandeel, et al. 2020), along with high divergence, complicates the estimation of synonymous divergence in SARS-CoV-2 and related viruses. In particular, in the third codon position the nucleotide frequency of T is 43.5% while it is just 15.7% for C. This resulting codon usage is not optimized for mammalian cells (e.g, (Chamary, et al. 2006)). A possible explanation is a strong mutational bias caused by Apolipoprotein B mRNA-editing enzymes (APOBECs) which can cause Cytosine-to-Uracil changes (Giorgio, et al. 2020).

A consequence of the skewed nucleotide frequencies is a high degree of homoplasy in synonymous sites that challenges estimates of *d*_*S*_. We here evaluated estimators of *d*_*S*_ in 300 bp sliding windows and found that a bias-corrected version of the maximum likelihood estimator tended to perform best for values of *d*_*S*_ < 2. We used this estimator to investigate the relationship between SARS-CoV-2 and related viruses in sliding windows. We show that synonymous mutations show shorter divergence to pangolin viruses, than the otherwise most closely related bat virus, RaTG13, in part of the receptor-binding domain of the *spike* protein. This strongly suggests that the previously reported amino acid similarity between pangolin viruses and SARS-CoV-2 is not due to convergent evolution, but more likely is due to recombination. In the recombination analysis, we identified recombination from pangolin strains into SARS-CoV-2, which provides further support for the recombination hypothesis. However, we also find that the synonymous divergence between SARS-CoV-2 and pangolin viruses in this region is relatively high, which is not consistent with a recent recombination between the two. It instead suggests that the recombination was into RaTG13 from an unknown strain, rather than between pangolin viruses and SARS-CoV-2, as proposed in (Boni, et al. 2020). The alternative explanation of recombination into SARS-CoV-2 from the pangolin virus, would require the additional assumption of a mutational hotspot to account for the high level of divergence in the region between SARS-CoV-2 and the donor pangolin viral genome. To fully distinguish between these hypotheses, additional strains would have to be discovered that either are candidates for introgression into RaTG13 or can break up the lineage in the phylogenetic tree between pangolin viruses and RaTG13.

The fact that synonymous divergence to the outgroups, RaTG13 and RmYN02, is not fully saturated, provides an opportunity for a number of different analyses. First, we can date the time of the divergence between the bat viruses and SARS-CoV-2 using synonymous mutations alone. In doing so, we find estimates of 51.71 years (95% C.I., 28.11-75.31) and 37.02 years (95% C.I., 18.19-55.85), respectively. Most of the uncertainty in these estimates comes from uncertainty in the estimate of the mutation rate reported for SARS-CoV-2. As more data is being produced for SARS-CoV-2, the estimate should become more precise and the confidence interval significantly narrowed. We note that the mutation rate we use here are estimated based on the entire genome, which may differ from that in non-recombination regions. To address this problem, we downloaded all the SARS-CoV-2 sequences that are available until 2020-08-17 from GISAID, and obtained an estimate of 1:0.81 for the ratio of mutation rates in the recombination and non-recombination regions, using the “GTRGAMMA” model implemented in the RAxML (Stamatakis 2014). Given the length ratio between the two partitions is 1:4, the difference between the partitions will cause a slight overestimate of the mutation rate by ∼5%, which is relatively small compared to the confidence intervals and the potential for other unknown sources of uncertainty. However, we warn that a residual cause of unmodeled statistical uncertainty is deviations from the molecular clock. Variation in the molecular clock could be modeled statistically (see e.g., (Drummond, et al. 2006) and (Lartillot, et al. 2016)), but the fact that synonymous mutations are mostly saturated for more divergent viruses that would be needed to train such models, is a challenge to such efforts. On the positive side, we note that the estimates of *d*_*S*_ given in Table 3 in general are highly compatible with a constant molecular clock. Boni et al. (Boni et al. 2020) obtained divergence time estimates similar to ours using a very different approach based on including more divergent sequences and applying a relaxed molecular clock. We see the two approaches as being complimentary. In the traditional relaxed molecular clock approach more divergent sequences are needed that may introduce more uncertainty due to various idiosyncrasies such as alignment errors. Furthermore, the relaxed molecular clock uses both synonymous and non-synonymous mutations and is, therefore, more susceptible to the effects of selection. Our approach allows us to focus on just the relevant in­group species and to use only synonymous mutations. The disadvantage is that we cannot accommodate a relaxed molecular clock. However, the fact that both approaches provide similar estimates is reassuring and suggests that neither idiosyncrasies of divergent sequences, natural selection, or deviations from a molecular clock has led to grossly misleading conclusions

Another advantage of estimation of synonymous and nonsynonymous rates in the outgroup lineage, is that it can provide estimates of the mutational load of the current pandemic. The *d*_*N*_*/d*_*S*_ ratio is almost 14 times larger in the circulating SARS-CoV-2 strains than in the outgroup lineage. While some of this difference could possibly be explained by positive selection acting at a higher rate after zoonotic transfer, it is perhaps more likely that a substantial proportion of segregating nonsynonymous mutations are deleterious, suggesting a very high and increasing mutation load in circulating SARS-CoV-2 strains.

## Supporting information

supplementary informaiton

## Acknowledgements

We are grateful to Dr. E.C Holmes for providing the genome sequence of RmYN02. We thank Dr. Sergei L Kosakovsky Pond for providing aligned sequences of SARS-CoV-2. We also thank Dr. Adi Stern for discussion. The research was funded by Koret-UC Berkeley-Tel Aviv University Initiative in Computational Biology and Bioinformatics to RN.

## Data Availability

The pangolin virus sequences, GD410721 and GX_P1E, were downloaded from GISAID with accession numbers EPI_ISL_410721 and EPI_ISL_410539, respectively, and RmYN02 sequence was provided by E. C. Holmes. All other sequences analyzed in this study were downloaded from either NCBI Genbank or National Microbiology Data Cente (NMDC). The accession codes for non-human sequences can be found in Supplementary Table 2 and the accession codes for human sequences can be found in Supplementary Table 3.

