## supplementary informaiton for "Synonymous mutations and the molecular evolution of SARS-Cov-2 origins"

for

**Supplementary Table 1.** Nucleotide composition at the first and second positions of codons across virus genomes.

| | Position 1 of codons | | | | Position 2 of codons | | | |
|---|---|---|---|---|---|---|---|---|
| **Accession** | **T** | **C** | **A** | **G** | **T** | **C** | **A** | **G** |
| GD410721 | 22.80% | 16.40% | 30.17% | 30.63% | 30.16% | 23.01% | 31.17% | 15.67% |
| GX_P1E | 22.46% | 16.84% | 30.17% | 30.54% | 30.23% | 23.03% | 30.92% | 15.82% |
| RaTG13 | 22.74% | 16.59% | 30.30% | 30.37% | 30.27% | 22.93% | 31.17% | 15.62% |
| RmYN02 | 22.93% | 16.46% | 30.15% | 30.46% | 30.41% | 22.87% | 30.86% | 15.87% |
| Wuhan-Hu-1 | 22.92% | 16.39% | 30.17% | 30.52% | 30.25% | 22.92% | 31.20% | 15.62% |

**Supplementary Table 2**. Accessions and sources of non-human coronavirus genome sequences used in this study.

| Genome Name | Source | Accession ID |
|---|---|---|
| SARS coronavirus PUMC03, complete genome | GenBank | AY357076.1 |
| SARS coronavirus TW9, complete genome | GenBank | AY502932.1 |
| SARS coronavirus Sin3408L, complete genome | GenBank | AY559097.1 |
| BetaCoV/pangolin/Guangdong/P1L/2019 | Lam et al. (2020) | N/A |
| hCoV-19/pangolin/Guangxi/P1E/2017\|EPI_ISL_410539\|2017 | GISAID | EPI_ISL_410539 |
| hCoV-19/pangolin/Guangdong/1/2019\|EPI_ISL_410721\|2019 | GISAID | EPI_ISL_410721 |
| Bat SARS CoV Rs672/2006, complete genome | GenBank | FJ588686.1 |
| Bat coronavirus BM48-31/BGR/2008, complete genome | GenBank | GU190215.1 |
| Bat coronavirus Cp/Yunnan2011, complete genome | GenBank | JX993988.1 |
| SARS-related bat coronavirus isolate Longquan-140 orf1ab polyprotein, spike glycoprotein, envelope protein, membrane protein, and nucleocapsid protein genes, complete cds | GenBank | KF294457.1 |
| Rhinolophus affinis coronavirus isolate LYRa11, complete genome | GenBank | KF569996.1 |
| BtRs-BetaCoV/HuB2013, complete genome | GenBank | KJ473814.1 |
| BtRs-BetaCoV/YN2013, complete genome | GenBank | KJ473816.1 |
| Rhinolophus bat coronavirus BtCoV/4991 RNA-dependent RNA polymerase (RdRp) gene, partial cds | GenBank | KP876546.1 |

| | | |
|---|---|---|
| Bat coronavirus isolate MLHJC35 P1ab gene, partial cds | GenBank | KU182963.1 |
| Severe acute respiratory syndrome-related coronavirus strain BtKY72, complete genome | GenBank | KY352407.1 |
| Bat SARS-like coronavirus isolate Rs4081, complete genome | GenBank | KY417143.1 |
| Bat SARS-like coronavirus isolate Rs4084, complete genome | GenBank | KY417144.1 |
| Bat SARS-like coronavirus isolate Rf4092, complete genome | GenBank | KY417145.1 |
| Bat SARS-like coronavirus isolate Rs4255, complete genome | GenBank | KY417149.1 |
| Bat SARS-like coronavirus isolate Rs4874, complete genome | GenBank | KY417150.1 |
| Bat SARS-like coronavirus isolate Rs7327, complete genome | GenBank | KY417151.1 |
| Bat coronavirus isolate Anlong-103, complete genome | GenBank | KY770858.1 |
| Bat coronavirus isolate Anlong-112, complete genome | GenBank | KY770859.1 |
| Bat coronavirus strain 16BO133, complete genome | GenBank | KY938558.1 |
| Bat coronavirus Rc-CoV-3 S gene for Spike protein, partial cds | GenBank | LC469301.1 |
| Bat SARS-like coronavirus isolate bat-SL-CoVZC45, complete genome | GenBank | MG772933.1 |
| Bat SARS-like coronavirus isolate bat-SL-CoVZXC21, complete genome | GenBank | MG772934.1 |
| SARS coronavirus Urbani isolate icSARS-C7-MA, complete genome | GenBank | MK062184.1 |
| Coronavirus BtRl-BetaCoV/SC2018, complete genome | GenBank | MK211374.1 |
| Coronavirus BtRs-BetaCoV/YN2018A, complete genome | GenBank | MK211375.1 |
| Coronavirus BtRs-BetaCoV/YN2018B, complete genome | GenBank | MK211376.1 |
| Coronavirus BtRs-BetaCoV/YN2018C, complete genome | GenBank | MK211377.1 |
| Coronavirus BtRs-BetaCoV/YN2018D, complete genome | GenBank | MK211378.1 |
| Bat coronavirus RaTG13, complete genome | GenBank | MN996532.1 |
| Severe acute respiratory syndrome coronavirus 2 isolate Wuhan-Hu-1, complete genome | GenBank | NC_045512.2 |
| Pangolin coronavirus isolate MP789 genomic sequence | GenBank | MT084071.1 |

**Supplementary Table 3**. Accessions and sources of the human SARS-CoV-2 genome sequences used in this study.

| Genome Name | Source | Accession ID |
|---|---|---|
| BetaCoV/Wuhan/IPBCAMS-WH-01/2019 | GenBank | MT019529 |
| BetaCoV/Wuhan/WH-01/2019 | GenBank | LR757998 |
| WIV02 | GenBank | MN996527 |
| WIV04 | GenBank | MN996528 |
| WIV05 | GenBank | MN996529 |
| WIV06 | GenBank | MN996530 |
| WIV07 | GenBank | MN996531 |
| BetaCoV/Wuhan/IPBCAMS-WH-02/2019 | GenBank | MT019530 |
| BetaCoV/Wuhan/IPBCAMS-WH-03/2019 | GenBank | MT019531 |
| BetaCoV/Wuhan/IPBCAMS-WH-04/2019 | GenBank | MT019532 |
| BetaCoV/Wuhan/WH19008/2019 | NMDC | NMDC60013002_06 |
| BetaCoV/Wuhan/WH19005/2019 | NMDC | NMDC60013002_10 |
| 2019-nCoV_HKU-SZ-002a_2020 | GenBank | MN938384 |
| 2019-nCoV_HKU-SZ-005b_2020 | GenBank | MN975262 |
| BetaCoV/Korea/SNU01/2020 | GenBank | MT039890 |
| BetaCoV/Wuhan/WH-03/2019 | GenBank | LR757996 |
| BetaCoV/Wuhan/IPBCAMS-WH-05/2020 | GenBank | MT019533 |
| BetaCoV/Wuhan/WH19004/2020 | NMDC | NMDC60013002_09 |
| 2019-nCoV WHU01 | GenBank | MN988668 |
| BetaCoV/Wuhan/WH-04/2019 | GenBank | LR757995 |

| | | |
|---|---|---|
| BetaCoV/Wuhan/YS8011/2020 | NMDC | NMDC60013002_07 |
| SARS-CoV-2/WH-09/human/2020/CHN | GenBank | MT093631 |
| SARS0CoV-2/61-TW/human/2020/NPL | GenBank | MT072688 |
| SARS-CoV-2/Yunnan-01/human/2020/CHN | GenBank | MT049951 |
| 2019-nCoV/USA-WA1/2020 | GenBank | MN985325 |
| BetaCoV/Hangzhou/HZ-1/2020 20cov-1L | GenBank | MT039873 |
| 2019-nCoV/USA-CA2/2020 | GenBank | MN994468 |
| 2019-nCoV/USA-AZ1/2020 | GenBank | MN997409 |
| 2019-nCoV/USA-CA1/2020 | GenBank | MN994467 |
| BetaCoV/Japan/AI/I-004/2020 | GenBank | LC521925 |
| BetaCoV/Australia/VIC01/2020 | GenBank | MT007544 |
| SARS-CoV-2/105/human/2020/CHN | GenBank | MT135041 |
| SARS-CoV-2/CA6/human/2020/USA | GenBank | MT044258 |
| SARS-CoV-2/IL2/human/2020/USA | GenBank | MT044257 |
| SARS-CoV-2/233/human/2020/CHN | GenBank | MT135043 |
| 2019-nCoV/Japan/KY/V-029/2020 | GenBank | LC522972 |
| 2019-nCoV/Japan/TY/WK-012/2020 | GenBank | LC522973 |
| 2019-nCoV/USA-CA3/2020 | GenBank | MT027062 |
| 2019-nCoV/USA-CA5/2020 | GenBank | MT027064 |
| SARS-CoV-2/IQTC02/human/2020/CHN | GenBank | MT123291 |
| 2019-nCoV/Japan/TY/WK-501/2020 | GenBank | LC522974 |

| | | |
|---|---|---|
| 2019-nCoV/Japan/TY/WK-521/2020 | GenBank | LC522975 |
| 2019-nCoV/USA-WI1/2020 | GenBank | MT039887 |
| Taiwan/NTU01/2020 | GenBank | MT066175 |
| BetaCov/Taiwan/NTU02/2020 | GenBank | MT066176 |
| SARS-CoV-2/IQTC01/human/2020/CHN | GenBank | MT123290 |
| 2019-nCoV/USA-CA7/2020 | GenBank | MT106052 |
| SARS-CoV-2/01/human/2020/SWE | GenBank | MT093571 |
| SARS-CoV-2/Hu/DP/Kng/19-020 | GenBank | LC528232 |
| SARS-CoV-2/Hu/DP/Kng/19-027 | GenBank | LC528233 |
| 2019-nCoV/USA-CA8/2020 | GenBank | MT106053 |
| 2019-nCoV/USA-MA1/2020 | GenBank | MT039888 |
| 2019-nCoV/USA-TX1/2020 | GenBank | MT106054 |
| 2019-nCoV/USA-CA9/2020 | GenBank | MT118835 |
| SARS-CoV-2/IQTC04/human/2020/CHN | GenBank | MT123292 |
| SARS-CoV-2/IQTC03/human/2020/CHN | GenBank | MT123293 |
| SARS-CoV-2/SP02/human/2020/BRA | GenBank | MT126808 |
| SARS-CoV-2/WA2/human/2020/USA | GenBank | MT152824 |
| Wuhan-Hu-1 | GenBank | NC_045512 |

**Supplementary Table 4**. Regions of SARS-CoV-2 with unique best hits with other coronaviruses.

| Start Position | End Position | Accession | % Identity | Alignment length |
|---|---|---|---|---|
| 2721 | 2820 | KF294457.1 | 96.000 | 100 |
| 2731 | 2830 | KF294457.1 | 96.970 | 99 |

| | | | | |
|---|---|---|---|---|
| 2741 | 2840 | KF294457.1 | 96.970 | 99 |
| 2751 | 2850 | KF294457.1 | 96.629 | 89 |
| 29801 | 29900 | AY357076.1 | 100.000 | 100 |
| 29811 | 29910 | AY357076.1 | 100.000 | 93 |

**Supplementary Table 5.** Recombination detection along the virus sequences using multiple methods. The coordiates refer to the positions in the Wuhan-Hu-1 reference genome (converted from the alignment coordinates outputed from the RDP program).

| No. | Start | End | Recombinant Sequence(s) | Minor Parental Sequence(s) | Major Parental Sequence(s) | RDP | GENECONV | Bootscan | Maxchi | Chimaera | SiSscan | 3Seq |
|---|---|---|---|---|---|---|---|---|---|---|---|---|
| 1 | 21225 | 24252 | ^RmYN02 | Unknown (GX_P1E) | Wuhan-Hu-1 | 1.53E-118 | 2.33E-82 | 1.00E-120 | 3.10E-41 | 5.42E-14 | 1.36E-37 | 4.65E-140 |
| 2 | 22851 | 23094 | ^Wuhan-Hu-1 | GD410721 | RaTG13 | 2.74E-17 | NS | 9.10E-16 | 4.10E-06 | 1.21E-06 | 3.04E-05 | 5.55E-15 |
| 3 | 25965 | 28297 | ^RmYN02 | Unknown (RaTG13) | Wuhan-Hu-1 | 5.87E-33 | 7.04E-41 | 1.43E-42 | 3.35E-02 | 1.12E-18 | 1.03E-39 | 6.57E-10 |
| 4 | 26002* | 27859 | ^RmYN02 | RaTG13, Wuhan-Hu-1 | Unknown (GD410721) | 5.41E-06 | NS | 1.83E-02 | 1.30E-14 | 1.62E-12 | NS | 4.66E-13 |
| 5 | 14611 | 15225 | ^GD410721 | Unknown (GX_P1E) | RmYN02 | 1.57E-02 | NS | NS | 2.34E-05 | 1.28E-04 | NS | 6.37E-05 |
| 6 | 21812 | 22473 | ^GD410721 | Unknown (GX_P1E) | RaTG13 | 1.51E-25 | NS | 1.33E-21 | 1.49E-09 | 1.69E-03 | 5.56E-16 | 2.41E-03 |
| 7 | 21563 | 21800* | ^GD410721 | Unknown (GX_P1E) | Wuhan-Hu-1, RaTG13 | 6.48E-05 | NS | 7.20E-06 | NS | NS | NS | 1.19E-06 |
| 8 | 21225* | 21377* | ^RmYN02 | RaTG13 | Unknown (Wuhan-Hu-1) | NS | 1.63E-02 | 2.20E-04 | NS | NS | NS | 2.42E-02 |
| 9 | 23535 | 23636 | ^GX_P1E | Unknown (RmYN02) | Wuhan-Hu-1, RaTG13 | 5.11E-04 | 4.20E-03 | NS | NS | NS | NS | 9.64E-03 |

* The actual breakpoint position is undetermined (it was most likely either overprinted by a subsequent recombination event or off the edges of the analysed sequence fragments).

^ The recombinant sequence may have been misidentified (one of the identified parents might be the recombinant)

Minor Parent: Parent contributing the smaller fraction of sequence.

Major Parent: Parent contributing the larger fraction of sequence.

Unknown: Only one parent and a recombinant need be in the alignment for a recombination event to be detectable. The sequence listed as unknown was used to infer the existence of a missing parental sequence.

NS: No significant P-value was recorded for this recombination event using the particular method in question.

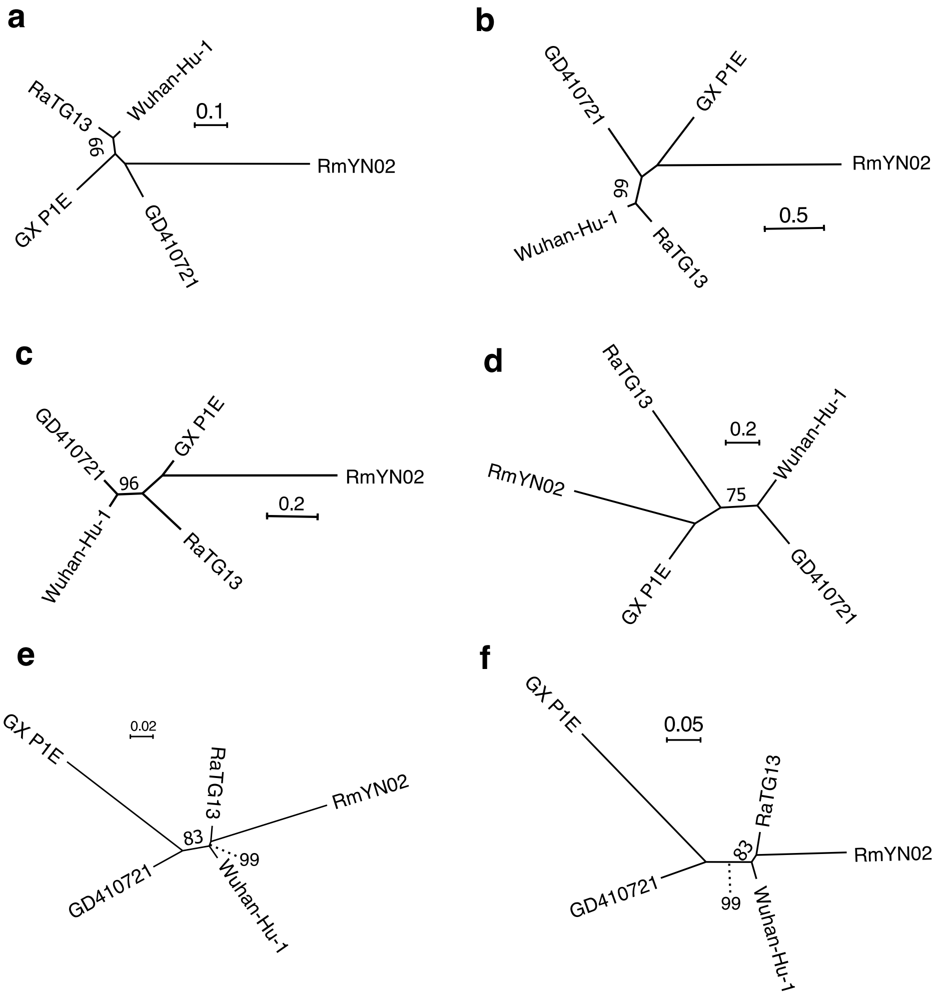


Supplementary Figure 1. Maximum-likelihood phylogenetic trees in the recombination regions. (a, c, e) phylogenetic trees constructed using full sequences in the recombination region 1-3 (Supplementary Table 5). (b, d, f) phylogenetic trees constructed by concatenating the 3rd bases of each codon in the recombination region 1-3. The bootstrap values are based on 1,000 replicates, only bootstrap values >= 70 are shown.

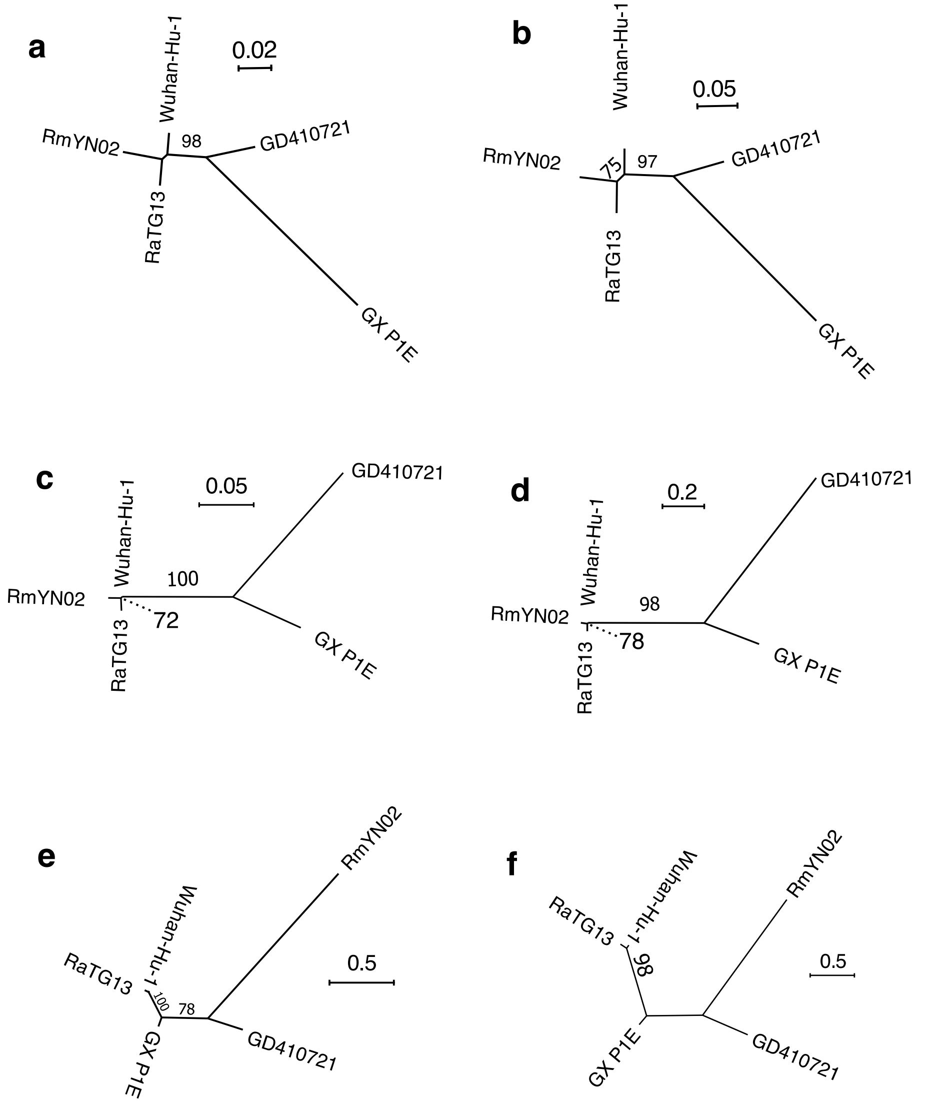


Supplementary Figure 2. Maximum-likelihood phylogenetic trees in the recombination regions. (a, c, e) phylogenetic trees constructed using full sequences in the recombination region 4-6 (Supplementary Table 5). (b, d, f) phylogenetic trees constructed by concatenating the 3rd bases of each codon in the recombination region 4-6. The bootstrap values are based on 1,000 replicates, only bootstrap values >= 70 are shown.

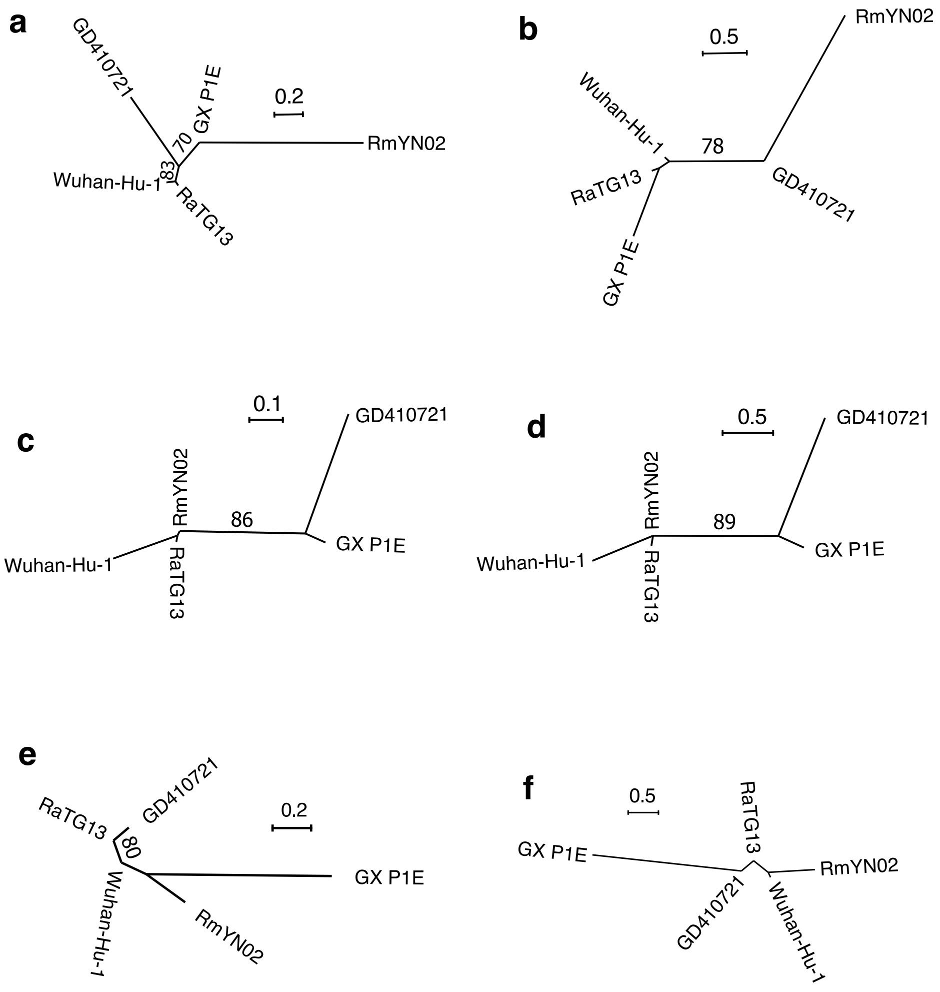


Supplementary Figure 3. Maximum-likelihood phylogenetic trees in the recombination regions. (a, c, e) phylogenetic trees constructed using full sequences in the recombination region 7-9 (Supplementary Table 5). (b, d, f) phylogenetic trees constructed by concatenating the 3rd bases of each codon in the recombination region 7-9. The bootstrap values are based on 1,000 replicates, only bootstrap values >= 70 are shown.
